# High efficiency multiplex biallelic heritable editing in *Arabidopsis* using an RNA virus

**DOI:** 10.1101/2022.01.20.477144

**Authors:** Ugrappa Nagalakshmi, Nathan Meier, Jau-Yi Liu, Daniel F. Voytas, Savithramma P Dinesh-Kumar

**Affiliations:** Department of Plant Biology, College of Biological Sciences, University of California, Davis, California, USA; The Genome Center, College of Biological Sciences, University of California, Davis, California, USA; Innovative Genomics Institute, University of California, Berkeley, CA, USA; Department of Genetics, Cell Biology and Development, University of Minnesota, St. Paul, Minnesota, USA; Center for Precision Plant Genomics, University of Minnesota, St. Paul, Minnesota, USA; Center for Genome Engineering, University of Minnesota, St. Paul, Minnesota, USA

**Keywords:** CRISPR, TRV, *Arabidopsis*, tRNA^Ileu^, agro-flooding, heritable gene editing, Flowering locus T, multiplex editing

## Abstract

Delivery of gene editing components such as the Cas nuclease and single guide RNAs (sgRNAs) into plant cells is commonly accomplished by agrobacterium-mediated transformation. Although *Arabidopsis* is easy to transform, generation of biallelic edited plants requires screening a large number of plants in subsequent generations. Here, we describe optimization of the *Tobacco rattle virus* (TRV) for *in planta* delivery of sgRNAs fused to a tRNA^Ileu^ that induces efficient multiplex somatic and biallelic heritable editing in *Arabidopsis*. Inclusion of tRNA^Ileu^ enhances the systemic movement of TRV and the mutant phenotype is visible in the initial TRV::sgRNA-tRNA^Ileu^ infected *Arabidopsis*, which allows for the uncovering of lethal phenotypes. Mutant progeny are recovered in the next generation (M1) at frequencies ranging from 30-60%, with 100% mutant recovery in the following (M2) generation. TRV::tRNA^Ileu^ system described here allows generation of biallelic edited plants in a single generation and is amenable for large-scale high throughput CRISPR screens.

## Introduction

Developing newer, more efficient gene editing methods in plants is critical to performing complex gene function studies and for the advancement of plant biotechnology. The advent and rapid deployment of CRISPR/Cas-based technology has allowed for the generation of mutant genotypes with highly specific mutations. CRISPR has been readily employed in plant biotechnology and adapted to many model and crop plant species^1,2,3^. So far, most methods for highly efficient, CRISPR-based gene editing rely on traditional transgenic approaches to deliver the Cas nucleases and single guide RNA (sgRNA) components. This is a major limitation since transgenes are left behind in the genome of transformed plants. This can lead to unknown off-target effects that can be difficult to mitigate.

As one of the first plant genomes to be sequenced, the model plant *Arabidopsis thaliana* has a plethora of genetic resources and decades of readily available experimental evidence from gene-function studies. However, one major limiting factor when using *Arabidopsis* for high throughput gene-function studies is the generation of stable, homozygous knockouts. Current, high-throughput, heritable editing technologies for testing gene function have not been adapted in *Arabidopsis*, as it relies on traditional agrobacterium-mediated transformation. While agrobacterium-mediated plant transformation in *Arabidopsis* can be a robust gene editing tool^4^, it suffers from several drawbacks. Namely, it requires several subsequent, self-crossing generations of large screening populations to generate biallelic homozygous mutant lines. Further, it leaves behind transgenes which could have unknown, off-target effects. Given the large-scale data that are being generated with omics-based approaches, it is increasingly important to be able to quickly evaluate the function of genes of interest to guide future research.

One option to deliver CRISPR editing components without the use of transgenics is to use viral vectors to deliver sgRNA targeted to a specific gene of interest in plants already expressing Cas nuclease^5,6^. Since the viral nucleic acid genome is not integrated into the host genome and is not transmissible to subsequent generations, progeny seeds of infected plants are virus and transgene free. Some advancements have been made in this area by engineering *Tobacco rattle virus* (TRV)^7,8^, *Potato virus X* (PVX)^9^, and *Barley stripe mosaic virus* (BSMV)^10^ viral vectors for delivery of sgRNAs targeted to genes of interest. A major limitation of using viral vectors is that many viruses are unable to enter meristematic tissue. As a result, gene editing is largely restricted from germline cells; hence, the gene editing is not heritable. Another major limitation of using viral vectors for gene editing is that each virus has a limited host range, so different viruses need to be developed and optimized for different plants.

Recently, we reported a novel method for delivery of sgRNAs that induces high efficiency somatic and heritable editing in *Nicotiana benthamiana* using a modified TRV vector system^8^. TRV is a single stranded positive sense RNA virus with a bipartite genome^11^. RNA1 of TRV encodes replicase proteins (134 kDa and a translational readthrough 194 kDa) from the genomic RNA, a 29 kDa movement protein (MP) from subgenomic RNA, and a 16 kDa cysteine-rich RNA silencing suppressor protein from a separate subgenomic RNA (Fig. 1a). RNA2 of TRV encodes the coat protein (CP) from the genomic RNA and two non-structural proteins (29.4 kDa and 32.8 kDa) from separate subgenomic RNAs (Fig. 1a). We engineered the TRV RNA2 by replacing the two non-structural proteins with multiple cloning sites (MCS) (Fig. 1a) or a gateway cloning cassette to facilitate virus-induced gene silencing (VIGS) to knockdown expression of target genes of interest in plants^12,13^. TRV has a wide host range and can infect many dicotyledonous plants. Additionally, TRV-VIGS vectors have been widely used by the scientific community for gene function studies in more than 25 plant species including *Arabidopsis*^11,14,15^.

**Fig. 1.**
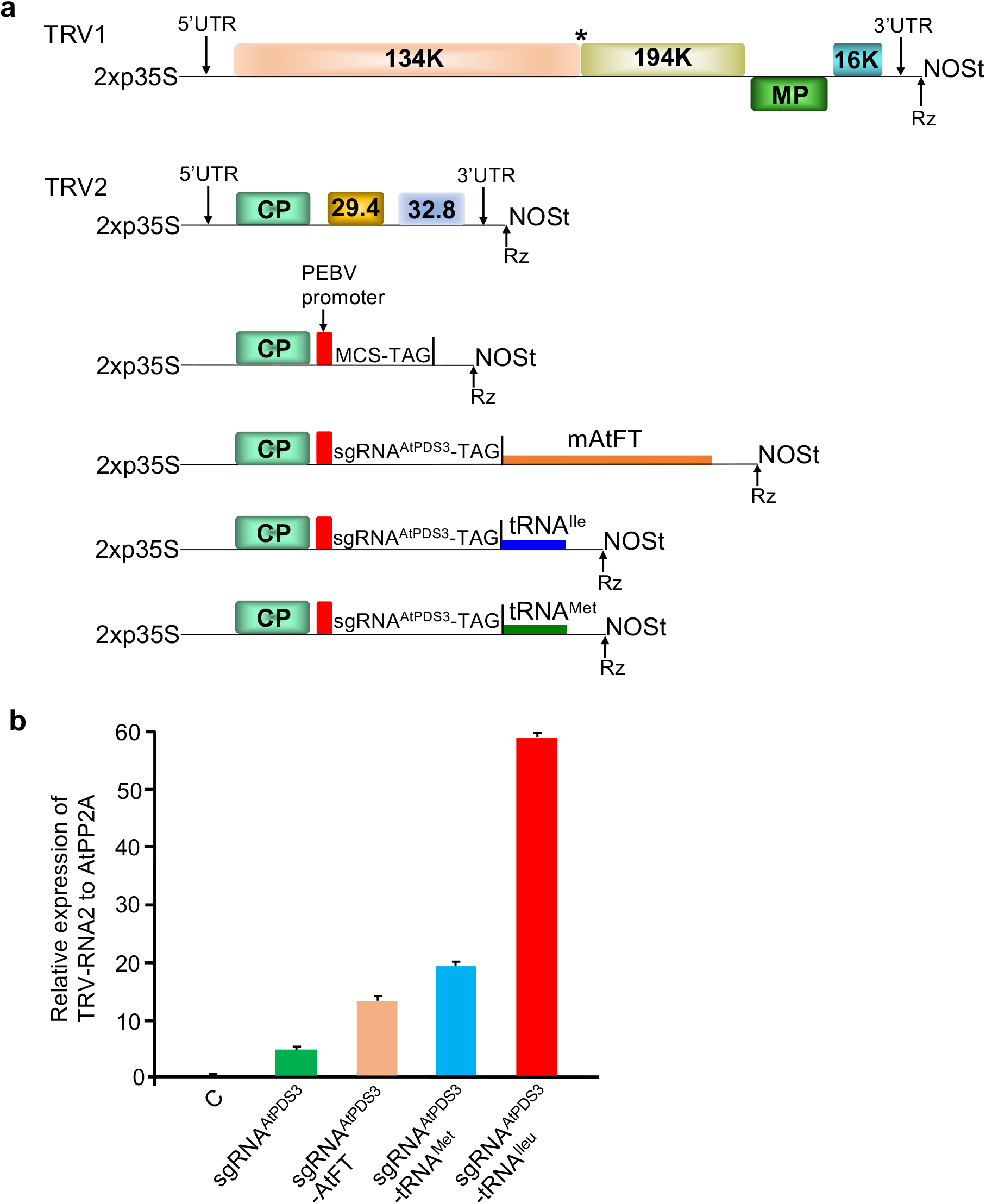
Inclusion of tRNA^Ileu^ in TRV promotes enhanced systemic movement. **a** Schematic representation of TRV1, TRV2 and various TRV2 derivative vectors. **b** Comparison of systemic movement of TRV with various mobile sequences. Relative levels of TRV RNA2 were measured by quantitative real-time PCR using *AtPP2A* as a reference. Data from three replicates were combined and values are shown as mean ±SD

A major advantage of TRV vectors is that they induce very mild symptoms on the infected plants^16,17^ and can transiently invade the meristematic region^18^. To express gene editing components such as sgRNAs, we modified the TRV RNA2 vectors used in VIGS by including a subgenomic promoter from another tobravirus, *Pea early browning virus* (PEBV) (Fig. 1a)^7^. Expression of sgRNAs using TRV with the pPEBV promoter (pPEBV) induced moderate editing of targeted genes in the infiltrated and systemic leaves of *N. benthamiana* plants^7^. However, the efficiency of heritable editing in the progeny plants was only 0.2%^7^. To overcome this, we recently reported a modified TRV system in which sgRNAs are expressed as fusion with mobile RNA sequences such as *Arabidopsis Flowering Locus T* (*AtFT*) and transfer RNAs (tRNAs) (Ellison et al., 2020). In *N. benthamiana*, TRV with sgRNAs augmented with mobile RNA sequences induced very high efficiency somatic editing in the infiltrated plants and 65-100% heritable editing efficiency in the next generation (M1)^8^. Recently, the expression of sgRNAs fused to *AtFT* sequence using PVX has also been shown to induce heritable editing with an efficiency of 22% in *N. benthamiana*^9^.

*N. benthamiana* serves as an excellent model species for viral studies due to the wide range of plant viruses able to infect it and ability to robustly express heterologous proteins^19,20^. However, it lacks the depth of genetic resources that are available for *Arabidopsis*. Hence, we sought to optimize our recently reported TRV system in *N. benthamiana* for use in *Arabidopsis*. While our studies were being conducted, another study showed that TRV with sgRNA flanked by self-splicing tRNA^21^ could induce low efficiency heritable epigene editing in the *Arabidopsis* M1 generation but not in the original infiltrated plants^22^. Furthermore, the geminivirus, *Cotton leaf crumple virus* (CLCrV) expressing a sgRNA has been shown to induce low efficiency heritable editing, but only in the second generation (M2) of *Arabidopsis* plants^23^. Here, we report a simple, optimized, agrobacterium-based flooding method combined with sgRNAs fused to the tRNA isoleucine (sgRNA-tRNA^Ileu^) that induces highly efficient multiplex somatic and heritable editing in *Arabidopsis*. The sgRNA-tRNA^Ileu^ is easily deliverable by TRV-based viral vector (TRV::sgRNA-tRNA^Ileu^). We show that the mutant phenotype was visible in the initial TRV::sgRNA-tRNA^Ileu^ infected plants and hence the system allows uncovering of lethal phenotypes. Furthermore, we observed 30-60% heritable editing efficiency in the M1 generation and 100% heritable editing efficiency in the M2 generation.

## Results

### Optimization of efficient delivery of sgRNA using TRV into *Arabidopsis* plants expressing *SpCas9*

We first tested TRV-based delivery of the sgRNA by using our previously established method for gene expression knockdown in *Arabidopsis* via VIGS^24^. We recently demonstrated that in *N. benthamiana*, TRV2 with a sgRNA fused to tRNA^Ileu^ moves more efficiently and generates high efficiency heritable editing compared to TRV with sgRNA alone^8^. Therefore, we aimed to adapt and optimize this system in *Arabidopsis*. We infiltrated TRV1 and TRV2 containing a sgRNA designed to target the *Arabidopsis thaliana PHYTOENE DESATURASE 3* (*AtPDS3*) gene fused to tRNA^Ileu^ (TRV2::sgRNA^AtPDS3^-tRNA^Ileu^) (Fig. 1a) into 2 week old *Arabidopsis* Col-0 plants expressing *SpCas9* (Col-0::*SpCas9*) (Supplementary Fig. 1a) by following the method described in^24^. About 3% of the infiltrated plants (1/36) showed visible photobleached spots or streaks on the leaves (Supplementary Fig. 2a). Since PDS in plants is required for the biosynthesis of photoprotective carotenoids, loss of function in *AtPDS3* will result in a photobleached phenotype.

Next, we tested two other methods to evaluate if somatic editing efficiency could be improved. In the first method, we grew plants on MS media for ten days and transferred the seedlings to a sterile petri plate with a mixture of agrobacterium harboring TRV1 and TRV2::sgRNA^AtPDS3^-tRNA^Ileu^ constructs. Then, needles were used to deliver multiple pricks to the leaves of individual plants (Supplementary Fig. 1b) (see Methods section for details). Twenty-four hours post-pricking, plants were transplanted into soil. In the second method, seeds were grown on MS plates for ten days, then flooded with a mixture of agrobacterium harboring TRV1 and TRV2::sgRNA^AtPDS3^-tRNA^Ileu^ (Supplementary Fig. 1c) (see Methods section for details). Three to four days post-flooding, plants were transplanted into soil. Only about 8% (3/36) of the pricked plants showed visible photobleached spots or streaks on the leaves. However, with the agrobacterium flooding method, about 22% of the plants (8/36) showed the photobleaching phenotype (Supplementary Fig. 2b). To assess that the observed phenotype is due to mutations in the *AtPDS3* gene, we PCR-amplified the region flanking the target site using DNA extracted from the entire leaves showing the photobleaching spots or streaks. Sanger sequencing of the PCR products followed by Synthego ICE (Inference of <u>C</u>RISPR <u>E</u>dits) analysis (https://ice.synthego.com/#/)^25^ confirmed the mutations in the *AtPDS3* gene (Supplementary Fig. 3). Together, these results indicated that the agro-flooding method is more efficient for the delivery of sgRNAs into *Arabidopsis* expressing *SpCas9* to induce editing.

### TRV with sgRNA^AtPDS3^ fused to tRNA^Ileu^ exhibits better systemic movement in *Arabidopsis* compared to sgRNA^AtPDS3^ fused to other mobile elements

Since the agro-flooding approach was more efficient, we followed this method in the rest of our experiments described in this manuscript. We recently reported that in addition to sgRNAs fused to tRNA^Ileu^, sgRNAs fused to *AtFT* without the start codon (m*AtFT*) and tRNA methionine (tRNA^Met^) also facilitate efficient movement of TRV in *N. benthamiana*^8^. Therefore, we tested systemic movement efficiency of TRV with sgRNA^AtPDS3^ fused to *mAtFT* (sgRNA^AtPDS3^-*mAtFT*) and tRNA^Met^ (sgRNA^AtPDS3^-tRNA^Met^) along with sgRNA^AtPDS3^-tRNA^Ileu^ (Fig. 1a). Ten day old Col-0 or Col-0::*SpCas9 Arabidopsis* seedlings on MS plates were flooded with agrobacterium harboring TRV1 and TRV2 with various sgRNA^AtPDS3^ mobile RNA fusions. After three to four days, seedlings were transplanted into soil. Three weeks after transplanting, the newly developed leaves were used for qRT-PCR analysis using primers that anneal to the 3’ end of TRV RNA2. A significantly higher amount of TRV was present in systemic leaves of plants infected with TRV2 with sgRNA^AtPDS3^-tRNA^Ileu^ compared to TRV2 with sgRNA^AtPDS3^ fused to other mobile RNA sequences, or when compared to sgRNA^AtPDS3^ with no mobile RNA fusion as a control (Fig. 1b). Systemic movement of TRV2 with sgRNA^AtPDS3^-tRNA^Met^ was better than that of sgRNA^AtPDS3^-*mAtFT* or sgRNA^AtPDS3^ alone (Fig. 1b). These results indicate that the sgRNA augmented with tRNA^Ileu^ moves better in *Arabidopsis* compared to other tested sequences.

### Efficient heritable editing of *AtPDS3* using TRV with sgRNA^AtPDS3^ fused to tRNA^Ileu^

Next, we set out to systematically characterize somatic editing efficiency of the *AtPDS3* gene using TRV2 with sgRNA^AtPDS3^-tRNA^Ileu^. Ten day old Col-0::*SpCas9* seedlings on MS plates were flooded with agrobacterium containing TRV1 and TRV2::sgRNA^AtPDS3^-tRNA^Ileu^ (Fig. 2a). Three days later, seedlings were transplanted into soil. About 15-20 days later, we observed the white photobleaching spots or streaks on the leaves (Fig. 2b). To determine the editing efficiency, we collected leaf tissue from completely photobleached regions, regions with overlapping photobleached and green (referred to as mosaic regions) and the green regions. DNA extracted from these samples was used to PCR amplify the region flanking the *AtPDS3* target site and the products were Sanger sequenced and subjected to Synthego ICE analysis. We observed a very high percentage of indels ranging from 60% to 99% at the *AtPDS3* target site in tissue collected from the completely photobleached regions (Fig. 2c and Fig. 3). In the mosaic regions, the editing efficiency varied from 20% to 70%, and editing efficiency was about 0% to 26% in the green regions (Fig. 2c and Fig. 3). During later stages of development, we observed complete photobleaching of some of the rosette and cauline leaves (Fig. 2d). In addition, some of the siliques and seed pods were completely photobleached (Fig. 2d and Supplementary Fig. 4). Together, these results indicate that sgRNA^AtPDS3^ fused to tRNA^Ileu^ induces mutations in the *AtPDS3* gene leading to the photobleached phenotype in *Arabidopsis*.

**Fig. 2.**
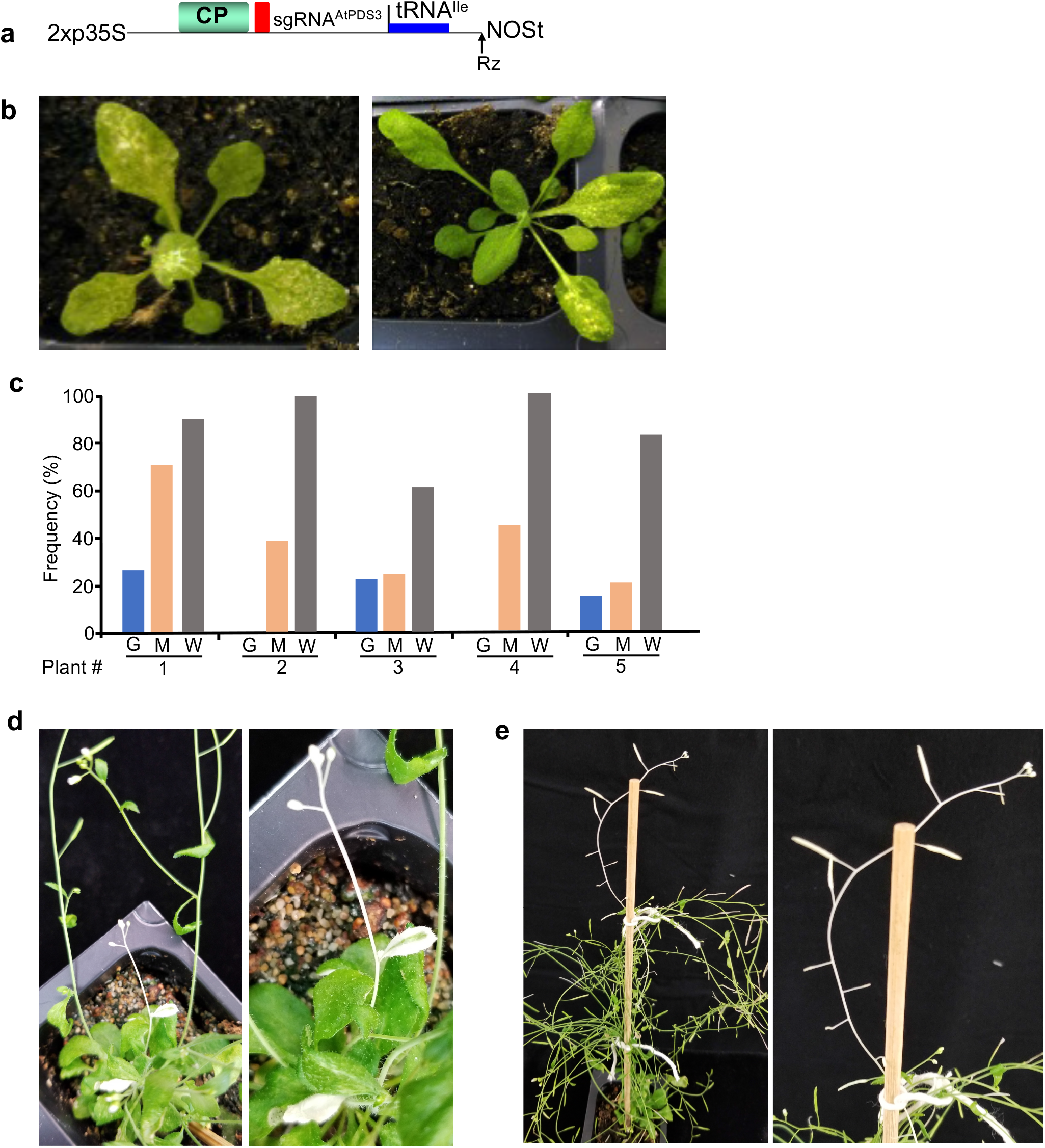
Efficient somatic editing in *AtPDS3* using TRV with tRNA^Ileu^. **a** Schematic of TRV RNA2 vector with sgRNA^AtPDS3^ fused to tRNA^Ileu^. **b** Phenotype of *Arabidopsis* plants infected with TRV1+TRV2::sgRNA^*AtPDS3*^-tRNA^Ileu^. White photobleached regions on the leaves is indicative of loss of *AtPDS3* function. **c** Editing efficiency in green (G), overlapping region between green and white (M) and white (W) region of leaves from 5 independent TRV1+TRV2::sgRNA^*AtPDS3*^-tRNA^Ileu^ infected *Arabidopsis* plants. Indel frequency was assessed by Sanger sequencing of amplicons spanning the *AtPDS3* target site followed by Synthego ICE analysis. **d** White photobleaching phenotype in some cauline leaves, stem and flowers due to editing in *AtPDS3*. Right panel is an enlarged version of part of the left panel. **e** White photobleaching phenotype of siliques. Right panel is an enlarged version of part of the left panel.

**Fig. 3.**
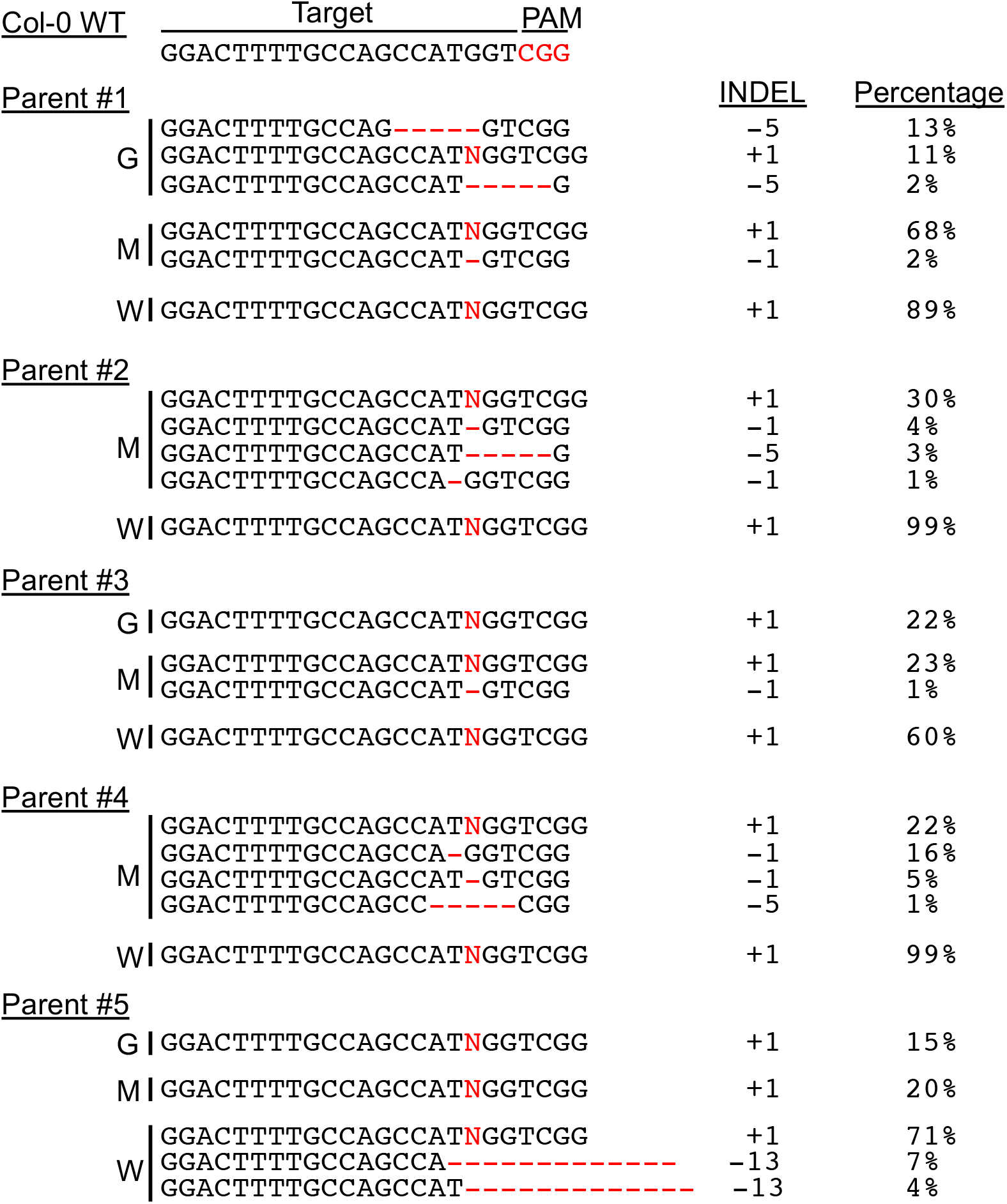
Mutation types and editing frequencies in *AtPDS3* in somatic tissue. The identified indel mutation types and indel frequencies derived from Sanger sequencing of amplicons spanning the *AtPDS3* target site followed by Synthego ICE analysis in green (G), overlapping region between green and white (M) and white (W) region of leaves from 5 independent TRV1+TRV2::sgRNA^*AtPDS3*^-tRNA^Ileu^ infected *Arabidopsis* plants shown in Fig. 2c.

Next, we assessed whether TRV2::sgRNA^AtPDS3^-tRNA^Ileu^ infected plants that show somatic editing also possess *AtPDS3* germline editing that is transmissible to the next generation. To accomplish this, we collected seeds from the aforementioned parent plants infected with TRV1 and TRV2::sgRNA^AtPDS3^-tRNA^Ileu^ (Fig. 2) in three batches between principal growth stage 8 to 5 days after principal growth stage 6.9^26^ that we refer to here as early, middle and late stage developed seeds. We observed very few photobleached seedlings collected from late stage developed seeds compared to early and middle stage developed seeds. From the middle stage developed seeds, 30-60% of progeny seedlings were completely photobleached, indicating biallelic mutations in *AtPDS3* (Fig. 4a and Supplementary Fig. 5). The next generation sequencing (NGS) of the *AtPDS3* PCR products flanking the target site (amplicons) of 8 white seedlings from parent 1 and parent 2 showed indel frequencies ranging from 86-93% and 95-99%, respectively (Fig. 4b and Supplementary Table 1). The NGS of amplicons of 8 green seedlings from each parent indicated indel frequency of 57-62% in these plants (Fig. 4b and Supplementary Table 2). We also performed Sanger sequencing of the region flanking the *AtPDS3* target region of 8 white progeny seedlings from two additional parents. Our Synthego ICE analysis indicated indel frequencies of 88-99% for progenies from parent 3 and 93-99% for progenies from parent 4 at the target site in *AtPDS3* (Supplementary Fig. 6). Together, these results indicate that TRV2::sgRNA^AtPDS3^-tRNA^Ileu^ induces high efficiency somatic and heritable editing in *Arabidopsis*.

**Fig. 4.**
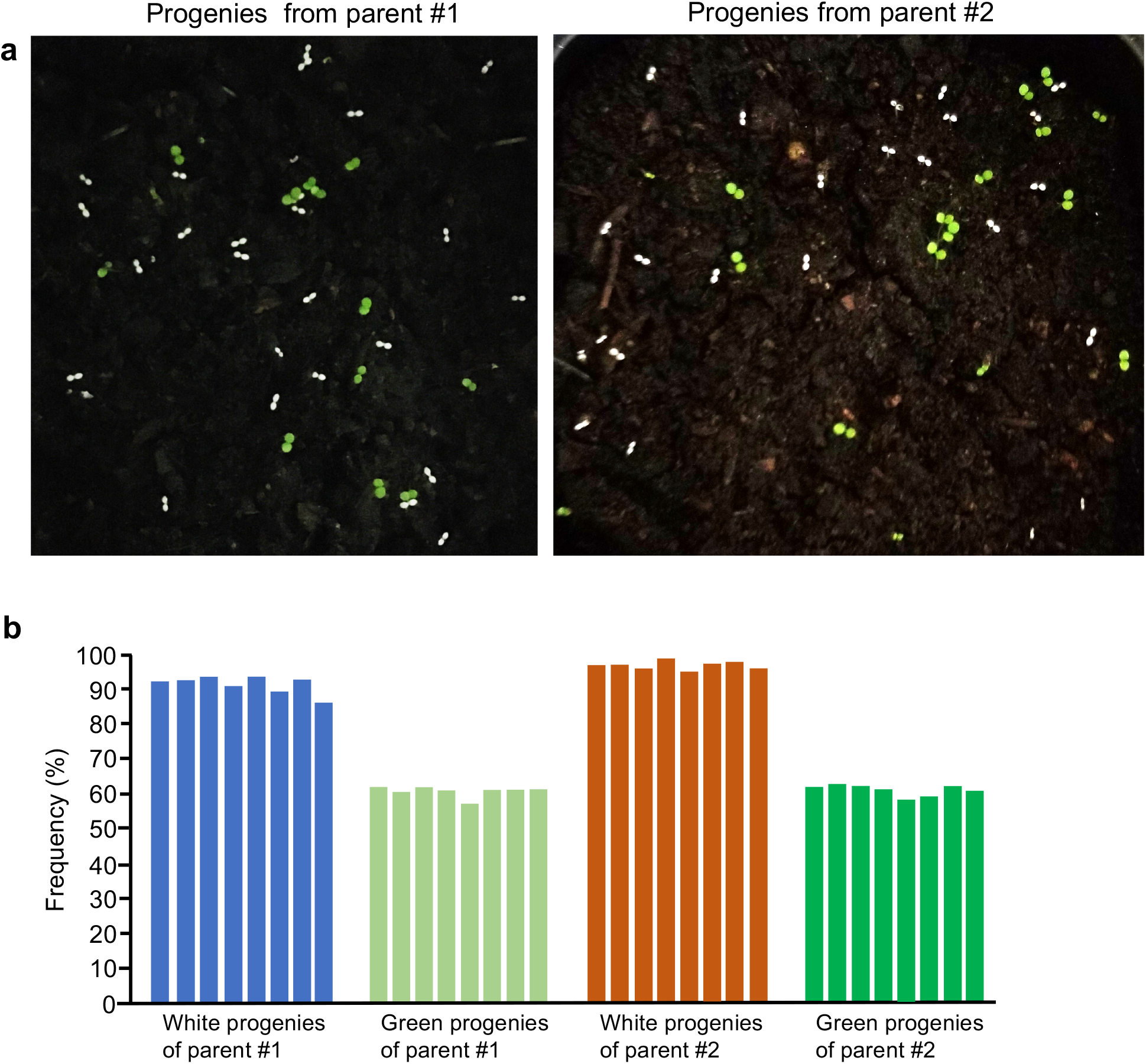
Heritable *AtPDS3* editing using TRV augmented with tRNA^Ileu^. **a** Phenotype of M1 progenies of seeds collected from two parental plants infected with TRV1+TRV2::sgRNA^AtPDS3^-tRNA^Ileu^. White photobleached seedlings are indicative of biallelic editing and loss of *AtPDS3* function. **b** Indel mutation frequencies in selected 8 white and 8 green progenies from each parent shown in (a) assessed by NGS of amplicons spanning the *AtPDS3* target site.

### Efficient multiplex editing of *AtCHLI1* and *AtCHLI2* using TRV with sgRNA fused to tRNA^Ileu^

To assess the efficiency of multiplex editing using this system, we targeted two genes that encode two isoforms of the I subunit of magnesium-chelatase (*CHLI*), *AtCHLI1* (At4g18480) and *AtCHLI2* (At5g45930). Along with CHLH and CHLD, CHLI is involved in catalyzing the first committed step toward chlorophyll synthesis. Knockout in both *AtCHLI1* and *AtCHLI2* genes results in albino sterile plants^27^. To target both *AtCHLI1* and *AtCHLI2* genes, we cloned two sgRNAs which target these genes in tandem with a 23-base pair (bp) spacer and placed them downstream of the PEBV promoter in TRV2 (TRV2::sgRNA^AtCHLI1^-tRNA^Ileu^-S-sgRNA^AtCHLI2^-tRNA^Ileu^) (Fig. 5a). Recently, we showed that the sgRNAs arranged in tandem with a 23 bp spacer works efficiently for multiplexing in *N. benthamiana*^8^.

**Fig. 5.**
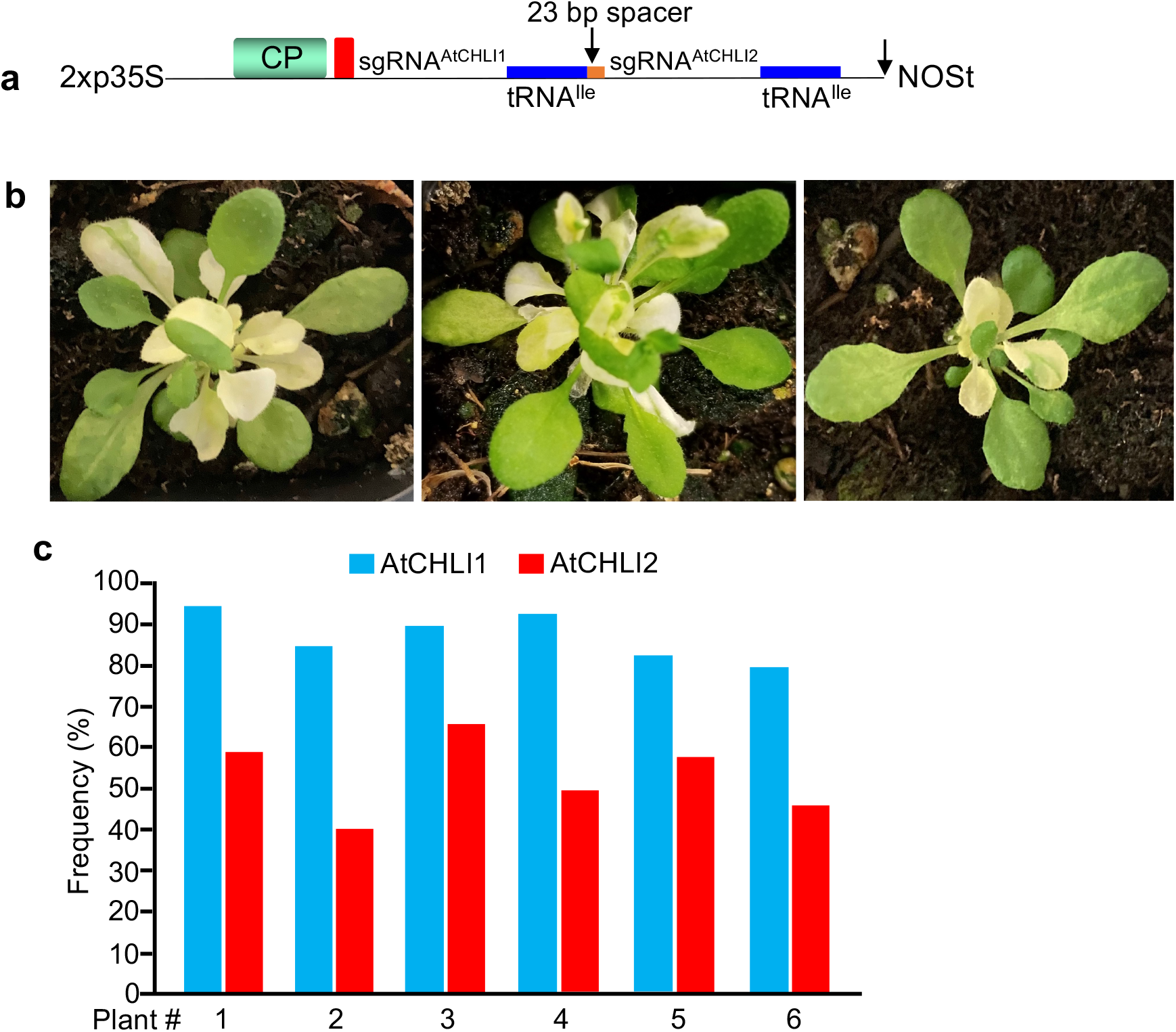
Efficient multiplex somatic editing in *AtCHLI1* and *AtCHLI2* using TRV augmented with tRNA^Ileu^. **a** Schematic of TRV RNA2 vector with sgRNA^AtCHLI1^ and sgRNA^AtCHLI2^ fused to tRNA^Ileu^ with a 23 bp spacer. **b** Phenotype of *Arabidopsis* plants infected with TRV1+TRV2::sgRNA^AtCHLI1^-tRNA^Ileu^-23bp-sgRNA^AtCHLI2^-tRNA^Ileu^. White albino and yellow leaves are indicative of editing and loss of function of both *AtCHLI2* and *AtCHLI2*. **c** Indel mutation frequencies in *AtCHLI1* and *AtCHLI2* from six independent plants showing albino and yellow leaves phenotype.

We performed a flooding treatment of ten day old Col-0::*SpCas9 Arabidopsis* seedlings with agrobacterium harboring TRV1 and TRV2::sgRNA^AtCHLI1^-tRNA^Ileu^-S-sgRNA^AtCHLI2^-tRNA^Ileu^. Starting from 15-20 days post-transplanting after TRV treatment, plants exhibited a yellow or white leaf color phenotype (Fig. 5b and Supplementary Fig. 7). The NGS amplicon sequencing of PCR products flanking *AtCHLI1* and *AtCHLI2* generated from plant tissue detected indel frequencies of 79-94% in the *AtCHLI1* gene and 45-66% in the *AtCHLI2* gene (Fig. 5C and Supplementary Tables 3 and 4). Together, these results indicate that TRV augmented with tRNA^Ileu^ can be used for multiplex gene editing in *Arabidopsis*. We were unable to estimate the heritable transmission of the knockout phenotype in the next generation because the parent plants showing phenotypes produced less seeds and the progeny resulting from these seeds were green or some were dead. We reasoned that since knocking out of both *AtCHLI1* and *AtCHLI2* genes results in sterility, the seeds that were produced carried wildtype genes.

### Efficient heritable multiplex editing of *AtTRY* and *AtCPC* genes using TRV with sgRNA fused to tRNA^Ileu^

Since we were unable to assess the heritability of multiplex editing in *AtCHLI1* and *AtCHLI2*, we targeted two MYB-like transcription factors *TRIPTYCHON* (*AtTRY*, At5g53200) and *CAPRICE* (*AtCPC*, At2g46410) using TRV2 augmented with tRNA^Ileu^. *AtTRY* and *AtCPC* functions as negative regulators of trichome development in *Arabidopsis*, since *try cpc* double mutant plants display increased trichomes on leaves leading to a clustered leaf trichomes phenotype^28,29^. We cloned an sgRNA fused tRNA^Ileu^ that targets both *AtTRY* and *AtCPC* under the control of the PEBV promoter in TRV2 (TRV2::sgRNA^AtTRY/AtCPC^-tRNA^Ileu^) (Fig. 6a). Agrobacterium harboring TRV1 and TRV2::sgRNA^AtTRY/AtCPC^-tRNA^Ileu^ was introduced into Col-0::*SpCas9 Arabidopsis* seedlings by the agro-flooding method as described above. Approximately 15-20 days post-transplanting of seedlings after agro-flooding we observed clustered trichomes on leaves (Fig. 6b and Supplementary Fig. 8). In addition to on the leaves, we also observed trichome clusters on the stems (Fig. 6b and Supplementary Fig. 8). To confirm that the observed clustered trichome phenotype is due to editing in the *AtTRY* and *AtCPC* genes, we PCR amplified the region flanking the *TRY* and *CPC* target sites using tissue from four independent plants and sequenced the amplicons by NGS. Our analysis indicated indel frequencies of 70-83% in the *AtCPC* gene and 71-76% in the *AtTRY* gene (Fig. 6c and Supplementary Tables 5 and 6). These results indicate that TRV with sgRNA^AtTRY/AtCPC^-tRNA^Ileu^ targets two genes simultaneously with high efficiency.

**Fig. 6.**
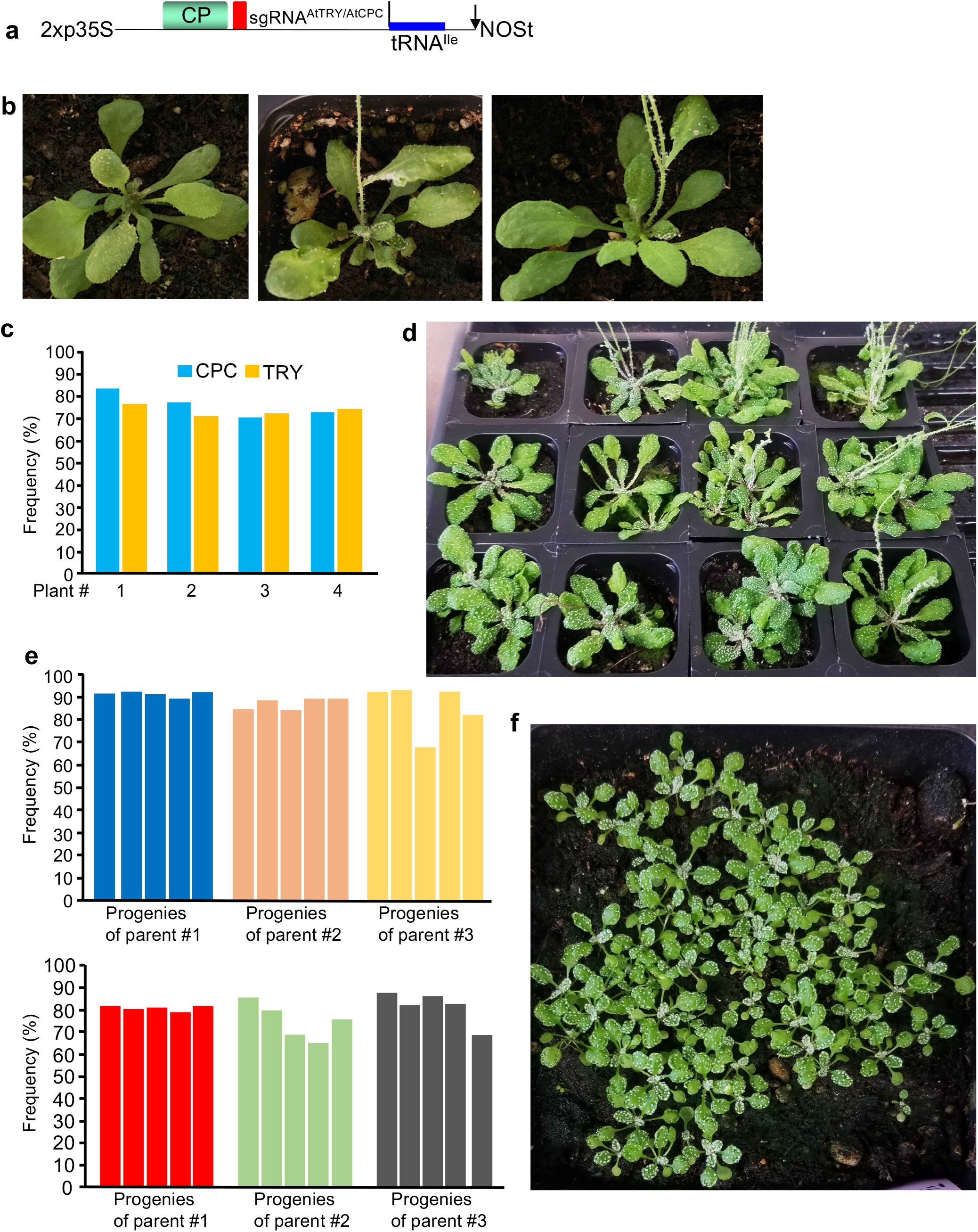
Efficient multiplex somatic and heritable editing in *AtTRY* and *AtCPC* using TRV augmented with tRNA^Ileu^. **a** Schematic of TRV RNA2 vector with single guide RNA targeting both *AtTRY* and *AtCPC* fused to tRNA^Ileu^. **b** Phenotype of *Arabidopsis* plants infected with TRV1+TRV2::sgRNA^AtTRY/AtCPC^-tRNA^Ileu^. Trichomes on the leaves and stems are due to editing and loss of function of both *AtTRY* and *AtCPC*. **c** Indel mutation frequencies in *AtTRY* and *AtCPC* from four independent plants showing leaf trichome phenotype. **d** Phenotype of M1 progenies of seeds collected from a parental plant infected with TRV1+TRV2::sgRNA^AtTRY/AtCPC^-tRNA^Ileu^. Trichomes on leaves and stems is indicative of biallelic editing in both *AtTRY* and *AtCPC*. **e** Indel mutation frequencies in five M1 progenies of three independent parent plants assessed by NGS of amplicons spanning the *AtCPC* (top panel) and *AtTRY* (bottom panel) target sites. **f** Phenotype of M2 progenies of seeds collected from a M1 progeny. All progenies showed leaf trichome phenotype indicating highly heritable biallelic editing in both *AtTRY* and *AtCPC*.

To assess the heritability of the observed editing in *AtTRY* and *AtCPC* in TRV1 and TRV2::sgRNA^AtTRY/AtCPC^-tRNA^Ileu^ infected plants, we collected seeds from parental plants that showed the clustered trichome phenotype (Fig. 6b and Supplementary Fig. 8) and grew them in soil. About 35-45% of the progeny plants from each parental line exhibited the clustered leaf trichomes phenotype (Fig. 6d and Supplementary Fig. 9a). The NGS amplicon sequencing of PCR products flanking the target sites indicated indel frequencies ranging from 89-92%, 84-89% and 68-93% in *AtCPC* gene (Fig. 6e, top panel and Supplementary Table 7) and 79-82%, 65-86% and 69-88% in *AtTRY* gene (Fig. 6e, bottom panel and Supplementary Table 8) in progenies from parent 1, parent 2 and parent 3 respectively.

To assess the transmission of knockout mutations in the 2^nd^ generation, we collected and planted the seeds from M1 plants that showed the clustered trichomes phenotype. We observed that 100% of the M2 progenies displayed the clustered leaf trichomes phenotype (Fig. 6f and Supplementary Fig. 9b). Together, these results indicate that TRV augmented with tRNA^Ileu^ induces highly efficient multiplex heritable knockout mutations in *Arabidopsis*.

## Discussion

Here, we describe an optimized TRV-based system for delivery of sgRNAs to induce highly efficient somatic and heritable editing in the model plant *Arabidopsis*. Our findings show that the fusion of tRNA^Ileu^ with sgRNA in TRV enhances systemic movement of the virus in *Arabidopsis* compared to sgRNA fused to *mAtFT*. This contrasts with our recent report which showed that both tRNA^Ileu^ and *mAtFT* promote better systemic movement of TRV in *N. benthamiana*^8^. Our findings are consistent with previous reports in *Arabidopsis* that the AtFT protein is the mobile form not the *AtFT* mRNA^30,31^. In wheat, systemic movement of BSMV with sgRNA fused to mutant wheat FT (BSMV-sg-mTaFT) was similar to the movement of BSMV expressing sgRNA alone (BSMV-sg)^10^. Although BSMV-sg-mTaFT induced similar somatic editing efficiency as BSMV-sg, BSMV-sg-mTaFT failed to induce heritable editing in wheat^10^. In rice, it was shown that mRNA of the FT ortholog, OsHd3a, was unable to move to the shoot apical meristem while the protein was fully mobile^32^. Due to the syntenic relationship between wheat and rice, this could be one reason why the BSMV-sg-mTaFT construct was unable to induce heritable editing.

Heritable editing in plants using viral vectors will depends on the ability of the virus to invade the meristematic region. Compared to most plant viruses, TRV can transiently invade the meristem early in the infection cycle^18^. Consistent with this, TRV has been used successfully to silence meristem genes like *NFL* (an ortholog of *Arabidopsis LEAFY*^16^) and floral homeotic genes such as *DEFICIENS* (an ortholog of *Arabidopsis AP3*^17^) in *N. benthamiana*. Despite this, TRV expressing sgRNA alone could only induce very low frequency heritable editing in *N. benthamiana*^7,8^. However, the addition of tRNA^Ileu^ to TRV significantly enhanced heritable editing frequency in *N. benthamiana*^8^ and *Arabidopsis* (this report). We hypothesize that enhanced viral systemic movement facilitated by tRNA^Ileu^ leads to more virus invading into the meristem region which, in turn, promotes editing in a higher numbers of germline cells. Future systematic analysis of TRV and TRV derivatives with mobile sequences should provide insights into engineering principles that would facilitate better invasion of TRV and other viruses into the meristem region.

Our findings in *Arabidopsis* (this report) and *N. benthamiana*^8^ show that the addition of tRNA^Ileu^ to sgRNAs in TRV enhances systemic movement of the virus and is capable of inducing efficient somatic and heritable editing in target genes. In a previous study, tRNA^Met^ and tRNA^Gly^ were identified as mobile tRNAs in the phloem exudates of pumpkin but not tRNA^Ileu 33^. When fused to a GUS transcript, tRNA^Met^ and tRNA^Gly^ facilitate unidirectional movement of GUS from shoot to root, but not from root to shoot in *Arabidopsis*^34^. This could be one reason for reduced infection rate of BSMV-tRNA^Met^ derivative in systemic leaves of wheat^10^. The addition of tRNA^Met^ into BSMV may enhance downward movement into roots than into the systemic leaves. In the case of TRV, our results showed that the inclusion of tRNA^Ileu^ significantly enhances the accumulation of TRV in the systemic leaves of *Arabidopsis* compared to tRNA^Met^. Future studies are required to understand the mechanistic basis of how tRNA^Ileu^ facilitates better movement of TRV and possibly other viruses in plants.

Recently, TRV has been shown to induce a low percentage of heritable epigene editing in *Arabidopsis*^22^. The epigene editing phenotype was only observed in the M1 progenies but not in the initial TRV inoculated *Arabidopsis* plants. We reason that the low heritability could be because TRV2 with self-splicing tRNA^21^ flanking the sgRNAs was used in these studies^22^. Since the tRNA will be spliced to generate sgRNAs, the TRV RNA2 would also be spliced reducing the infectiousness of the virus. In this scenario, there will be a population of TRV RNA2 population in which the tRNA is not spliced, this non-spliced tRNA will fail to produce sgRNAs with precise 5’ and 3’ ends. In BSMV, the inclusion of self-splicing tRNA flanking sgRNA significantly reduced the systemic virus infectivity in wheat^10^. The self-splicing tRNA sequence is very similar to the tRNA^Gly^, with the few base differences residing in the loop region of these tRNAs. TRV with sgRNA fused to tRNA^Gly^ did not enhance systemic movement of the virus and induced very low heritable gene editing in *N. benthamiana*^8^.

Very low efficiency, heritable editing observed only in the second (M2) generation in *Arabidopsis* has also been recently reported using the geminivirus CLCrV^23^. In that report, sgRNA fused to the N-terminus 102 bp of AtFT (AtFT^1-102bp^) was expressed using CLCrV. This study failed to observe mutant phenotypes in either the CLCrV-infected plants or in the M1 progenies, but a small number of mutant progenies in the M2 generation were recovered. Based on our findings in this paper and previous reports indicating AtFT mRNA is immobile in *Arabidopsis*^30,31^, we hypothesize that the very low efficiency of editing through CLCrV with AtFT^1-102bp^ that was observed could be due to the virus being unable to efficiently invade the meristem.

The TRV system with tRNA^Ileu^ that we describe in this paper is the most efficient method for inducing somatic and heritable editing in *Arabidopsis* reported to date. Furthermore, we show that the system is suitable for multiplex editing with very high efficiency. It also facilitates uncovering lethal phenotypes because one could observe the phenotype in the original virus infected plants. Our findings indicate that biallelic mutants can be generated in a single generation. In the second generation, 100% of the M2 progenies exhibited the mutant phenotype. Introducing gene editing components into *Arabidopsis* through transformation is relatively easy compared to other plants; however, it still requires a significant amount of time to generate biallelic knockout mutants using CRISPR/Cas9. The optimized and simple agro-flooding method combined with TRV augmented with tRNA^Ileu^ described here is amenable for high throughput editing of candidate genes identified from different omics approaches.

## Methods

### Vector construction

The pYL192 (TRV1^13^; Addgene ##148968), SPDK3876 (TRV2 with PEBV promoter^8^; Addgene #149275) have been described. TRV2-pPEBV-MCS-tRNA^Ileu^ (SPDK3888, Addgene #149276), TRV2-pPEBV-MCS-tRNA^Met^ (SPDK3889, Addgene #149277), and TRV2-pPEBV-MCS-mAtFT (SPDK3895) have been described^8^. To generate sgRNA^AtPDS3^ vectors, oligonucleotides 9208 and 9209 and 8788 and 8789 were annealed and ligated into *Xba*I-*Sac*I cut SPDK3888, SPDK3889, SPDK3895 and SPDK3876 resulting in SPDK3959 (TRV2-sgRNA^AtPDS3^-tRNA^Ileu^), SPDK3952 (TRV2-sgRNA^AtPDS3^-tRNA^Met^), SPDK3969 (TRV2-sgRNA^AtPDS3^-tRNA^mAtFT^), and SPDK4151 (TRV2-sgRNA^AtPDS3^) vectors respectively. To generate sgRNAs targeting *AtCHLI1* and *AtCHLI2* with 23 bp spacer (SPDK4146), PCR products generated using 100272 and 100149 primers with SPDK3959 DNA template and 8787 and 100273 primers with SPDK3959 DNA template and Phusion DNA polymerase were digested with *Xba*I-*Eag*I and *Eag*I-*Sac*I respectively and cloned into SPDK3888 cut with *Xba*I-*Sac*I. The vector containing sgRNA targeting *AtCPC* and *AtTRY* was generated by cloning PCR product generated using 100279 and 8787 primers with SPDK3959 template was digested with *Xba*I-*Sac*I and cloned into SPDK3888 cut with same enzymes. All primers used for cloning are listed in Supplementary Table 9.

### Plant growth conditions

Col-0, Col-0::*SpCas9* (line 13-19) or TRV infected plant progeny seeds were stratified at 4° C in water in the dark for 3 days before sowing in soil. Plants were grown in a growth chamber set at 26°/22°C day/night condition with a 12-h light/12-h dark photoperiod and a light intensity of 100 μE m^-2^ sec^-1^, 50% humidity. Col-0 and Col-0::*SpCas9* seedlings from MS plates treated with agrobacterium with TRV were transplanted to soil and grown at the same conditions as plants started from the seeds.

### Preparation of agrobacterium harboring TRV vectors

TRV1, TRV2 and various TRV2 derivative vectors were introduced into chemically competent agrobacterium strain GV3101. Transformants were confirmed by colony PCR. Agrobacterium harboring TRV vectors were grown in 5 ml of lysogeny broth (LB) with antibiotics for 16-18 h at 28° C. Agrobacterium cultures were centrifuged for 20 min at 3,500 relative centrifugal force (RCF). The LB medium was discarded and agrobacterium cells were resuspended in 5 ml of sterile water and centrifuged for 10 min 3,500 RPM. The resulting pellets were resuspended in sterile agro-infiltration buffer [10 mM MgCl2, 10 mM 2-(*N*-morpholino) ethanesulfonic acid, and 250 µM acetosyringone] to OD_600_ = 1.5 and incubated with slow shaking for 3 h at room temperature. The TRV1 and TRV2 or TRV2 derivatives were mixed at 1:1 ratio and used for various inoculation methods.

### TRV inoculation

#### Leaf infiltration

Agrobacterium with TRV1 and TRV2 or TRV2 derivatives were infiltrated into 2 weeks old Col-0::*SpCas9* plants using a 1mL needleless syringe as described in^24^. About 10 days post-infiltration, the infiltrated plants were monitored every day for any visible phenotypes.

#### Agro-pricking with a needle

Col-0 and Col-0::*SpCas9* seeds were surface sterilized by placing in a 1.5 mL sterile tube and 1 ml 75% ethanol and 0.1% Triton X-100 (Sigma-Aldrich) was added and placed in Eppendorf Mixer 5432 for 4-5 minutes. Seeds were washed with 1 ml of 100% ethanol 2 times with 3 minute incubation each time in the mixer and seeds were dried overnight in the laminar hood. Sterilized seeds were placed on one-half strength MS medium containing Gamborg vitamins (Phyto Technologies Laboratories) solidified with 0.25% Phytoblend (Caisson Labs) in petri plates (100 mm × 25 mm). The MS plates with seeds were covered in aluminum foil and kept for three days at 4°C and transferred to a growth chamber set at 24°C with a light intensity of 100 μE m^-2^ sec^-1^ and a 12 h light/12 h dark photoperiod. Eight to ten days post-germination, the seedlings were transferred to a sterile petri plate and 10 ml of agrobacterium suspension with TRV1 and TRV2 or TRV2 derivatives was added. Leaves of each seedling were pricked using Monoject® Hypodermic Plastic Hub Needle 25g x 1”. The seedlings were incubated overnight at room temperature and next day they were planted on soil and grown in a growth chamber as described above.

#### Agro-flooding

Col-0 and Col-0::*SpCas9* seeds were sterilized and set on MS plates as described above. Ten ml agrobacterium suspension with TRV1 and TRV2 or TRV2 derivatives was dispensed directly into the MS plates containing eight to ten days-old *Arabidopsis* seedlings. The plates were incubated for 4 days in the growth chamber at 24°C with a light intensity of 100 μE m^-2^ sec^-1^ and a 12 h light/12 h dark photoperiod. Seedlings were transplanted to soil and grown in a growth chamber as described above.

### Quantitative real-time PCR analysis

Leaf tissue was collected from control and TRV infected plants and total RNA was extracted using TRIzol reagent (Life Technologies). Total RNA was treated with RNase free DNase I (New England Biolabs). First-strand cDNA was synthesized using SuperScript III reverse transcriptase (Life Technologies) and TRV-specific primer SP9239 or the control AtPP2A-R primer. RT–qPCR was performed using the Bio-Rad CFX96 qPCR instrument (Bio-Rad) with iTaq Universal SYBR Green Supermix (Bio-Rad), 2.5 μM each of TRV-specific primers (9238 and 9239) or AtPP2a-specific primers (AtPP2A-F and AtPP2A-R) and 5% diluted first-strand cDNA. Data presented are normalized for AtPP2A expression level.

### Analysis of somatic and heritable editing efficiency in *Arabidopsis*

To assay editing in TRV infected plants or M1 progenies, Phire Plant Direct PCR Kit (Thermo Fisher Scientific) was used. A small leaf or a piece of leaf of about 2 mm in diameter was collected into a tube containing 20 μl of dilution buffer and the leaf sample was crushed using a 1 ml pipette tip. Centrifuged for 5 min at 14,000 rpm and the supernatant was transferred to a new tube. PCR reaction was performed using 2-5 μl of this genomic DNA as template and 25 pmol of primers in a 50 μl reaction. All primers used for genotyping are listed in Supplementary Table 9. The resulting amplicons were purified using Zymoclean Gel DNA Recovery Kit (Zymo Research). The amplicons were sent for Sanger sequencing (Eurofins Genomics) or Amplicon-Ez Next Generation Sequencing (NGS) (Genewiz/Azenta Life Sciences). Sanger trace sequencing file (.ab1 file) was used to assess indel percentage using Synthego ICE (Inference of CRISPR Edits) analysis pipeline (https://ice.synthego.com/) (Hsiau et al., 2019). The paired-end amplicon sequencing reads from Amplicon-EZ were de-multiplexed using ea-utils^35^. The demultiplexed reads were uploaded to Cas-Analyzer^36^ and compared against the targeted DNA sequence. In order to avoid the quantification of indels located far from the target sequence, we used a 40 bp window to either side of the target site for analysis. We also filtered out sequences with <5 reads to eliminate any sequencing errors from the analysis.

## Data availability

Data supporting the findings of this work are available within the paper and its Supplementary Information files, or from the corresponding author upon request.

## Competing Interests

The authors declare no competing interests.

## Acknowledgements

We thank Professor Jen Sheen and Dr. Jian-Feng Li for providing *SpCas9* expressing *Arabidopsis* line. We thank April DeMell for critical reading and editing the manuscript. This work was supported by Innovative Genomics Institute (to S.P.D-K and U.N), Agricultural Innovation through Gene Editing program grant no. 2020-67013-31544/project accession no. 1022332 from the USDA National Institute of Food and Agriculture (to S.P.D-K and U.N), and grant no. HR0011-17-2-0053 from the Defense Advanced Research Projects Agency (DARPA; to S.P.D-K and D.F.V).

## Author Contributions

U.N and S.P.D-K conceptualize the project and designed the experiments. U.N and J-Y.L performed the experiments. U.N and N.M analyzed the data and interpreted the results. D.F.V provided suggestions and discussed the results with S.P.D-K. Y.X. and J.S generated and characterized *SpCas9* expressing *Arabidopsis* transgenic line. U.N., N.M., and S.P.D-K wrote the original draft of the manuscript. All authors reviewed and edited the manuscript.

---

**Supplementary Fig. 1.**
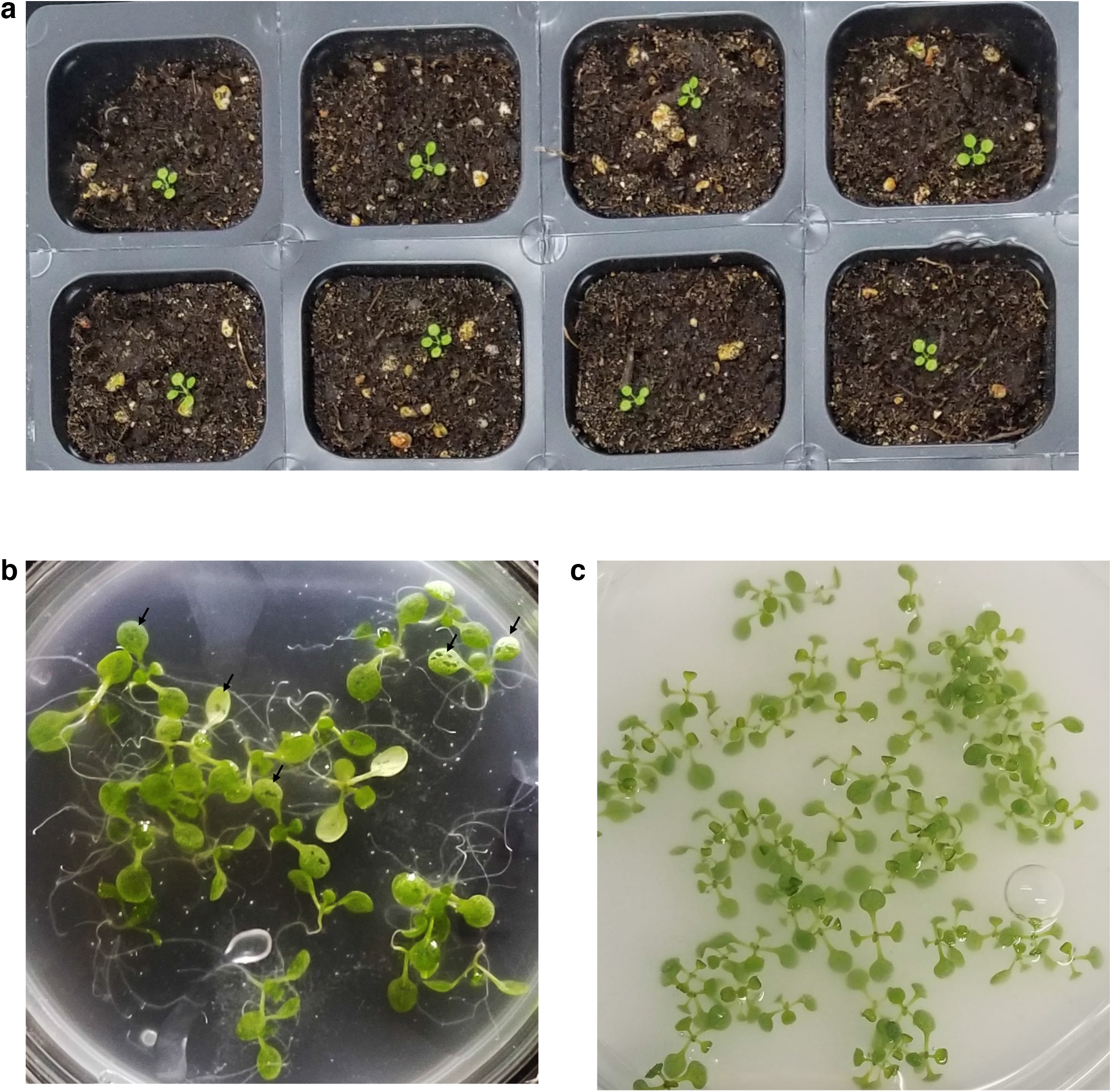
Different methods of introducing TRV vectors with sgRNAs into *Arabidopsis*. **a** Plants used for syringe infiltration of agrobacterium harboring TRV vectors. **b** Plate with seedlings in which leaves are pricked with agrobacterium harboring TRV vectors (black arrows). **c** Plate with seedlings flooded with agrobacterium harboring TRV vectors.

**Supplementary Fig. 2.**
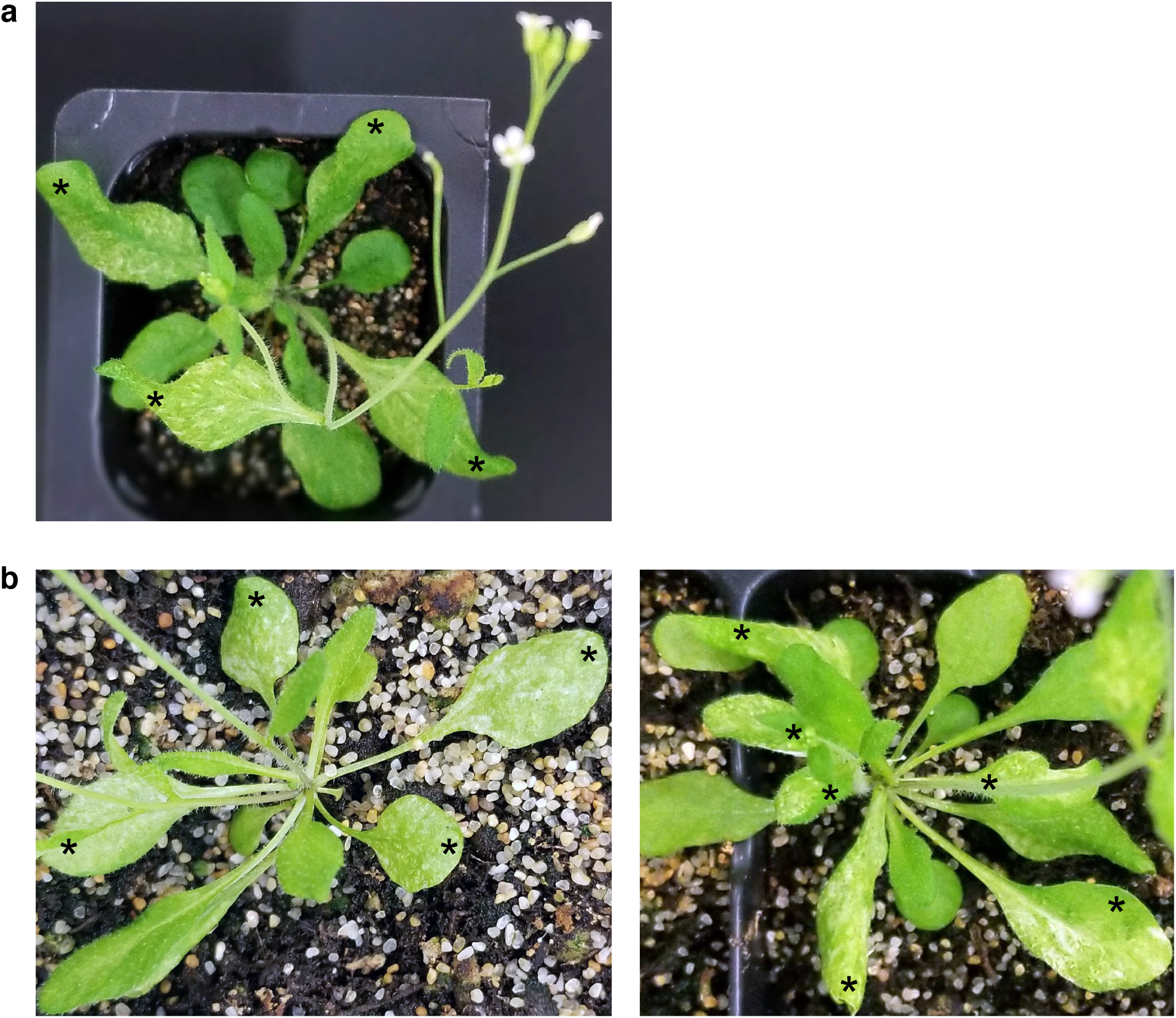
Somatic editing in *AtPDS3* using TRV augmented with tRNA^Ileu^. **a** Phenotype in systemic leaves of *Arabidopsis* plant that was syringe infiltrated with agrobacterium harboring TRV1 and TRV2::sgRNA^AtPDS3^-tRNA^Ileu^. White photobleached region on the leaves (asterisks) is due to editing and loss of AtPDS3 function. **b** Phenotype in systemic leaves of transplanted *Arabidopsis* plants that were treated by flooding of agrobacterium harboring TRV1 and TRV2::sgRNA^AtPDS3^-tRNA^Ileu^. White photobleached region on the leaves (asterisks) is due to editing and loss of AtPDS3 function.

**Supplementary Fig. 3.**
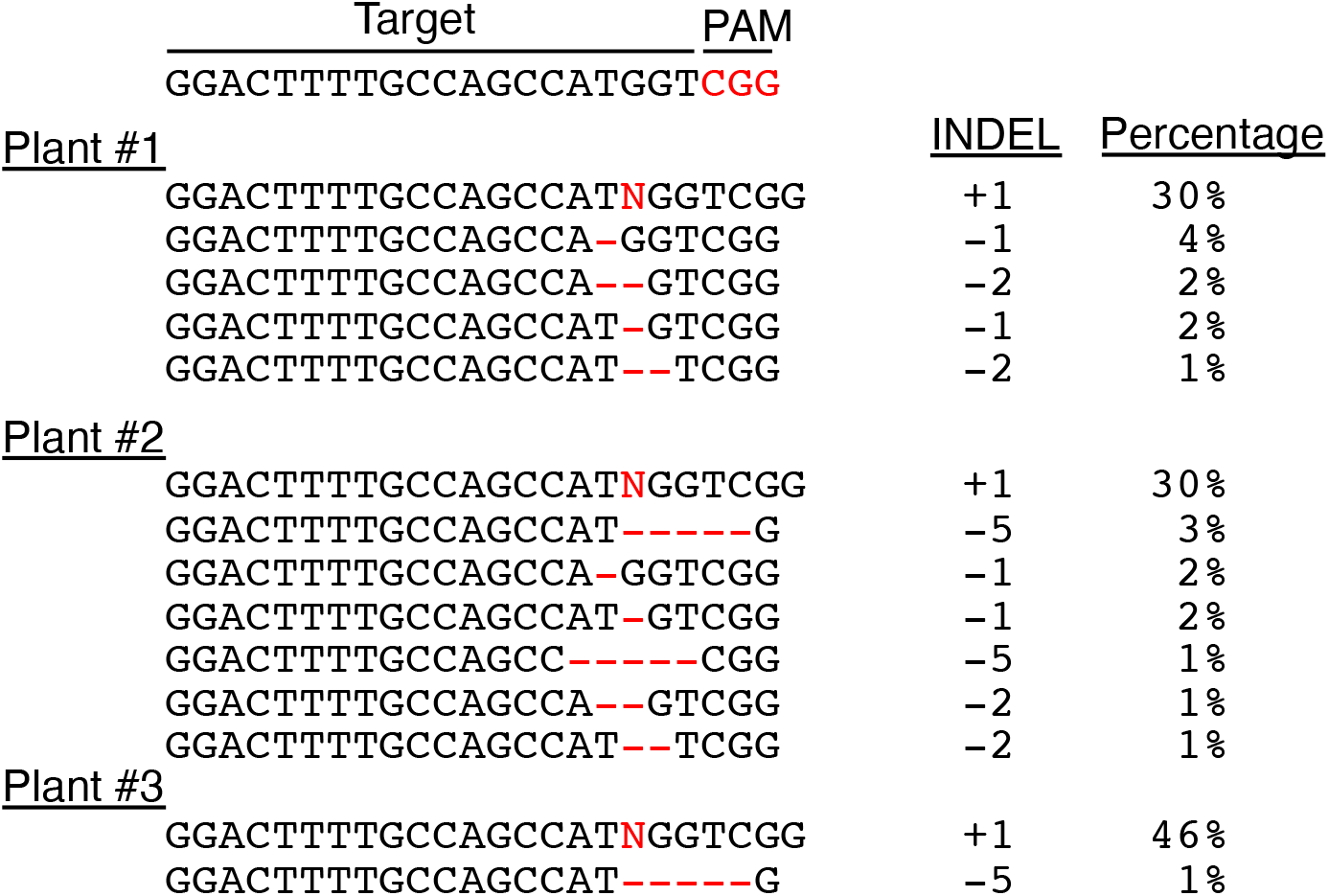
Indel mutation types in *AtPDS3* in leaves showing photobleaching phenotype. The indel mutation types and frequencies identified from Sanger sequencing of amplicons spanning the *AtPDS3* target site in leaves of three independent plants that were treated with agro-flooding and showed a photobleaching phenotype.

**Supplementary Fig. 4.**
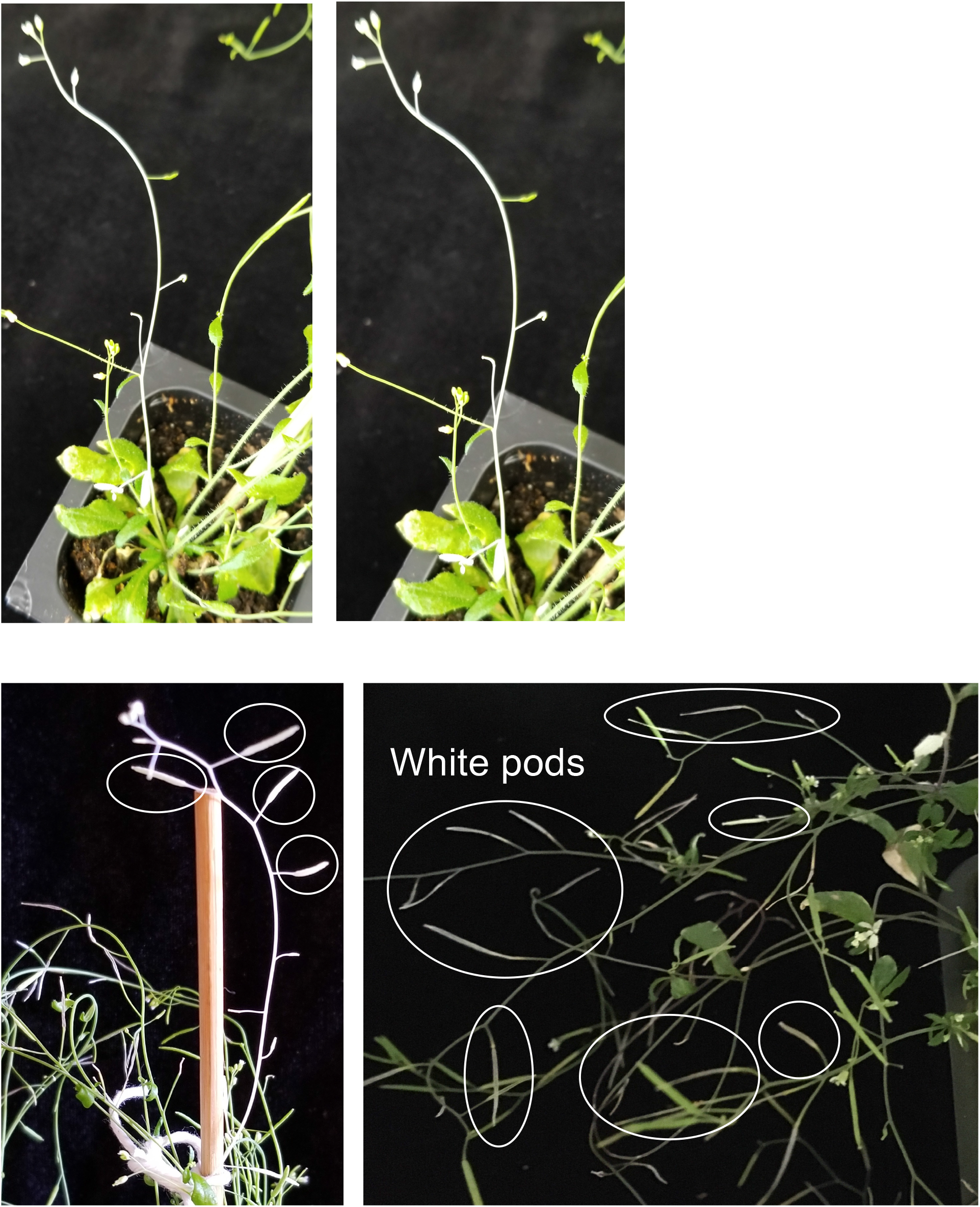
Photobleaching of stems, flowers and siliques of plants infected with TRV expressing SgRNA^AtPDS3^ fused to tRNA^Ileu^. Left panel of the top panel is an enlarged version of part of the right panel.

**Supplementary Fig. 5.**
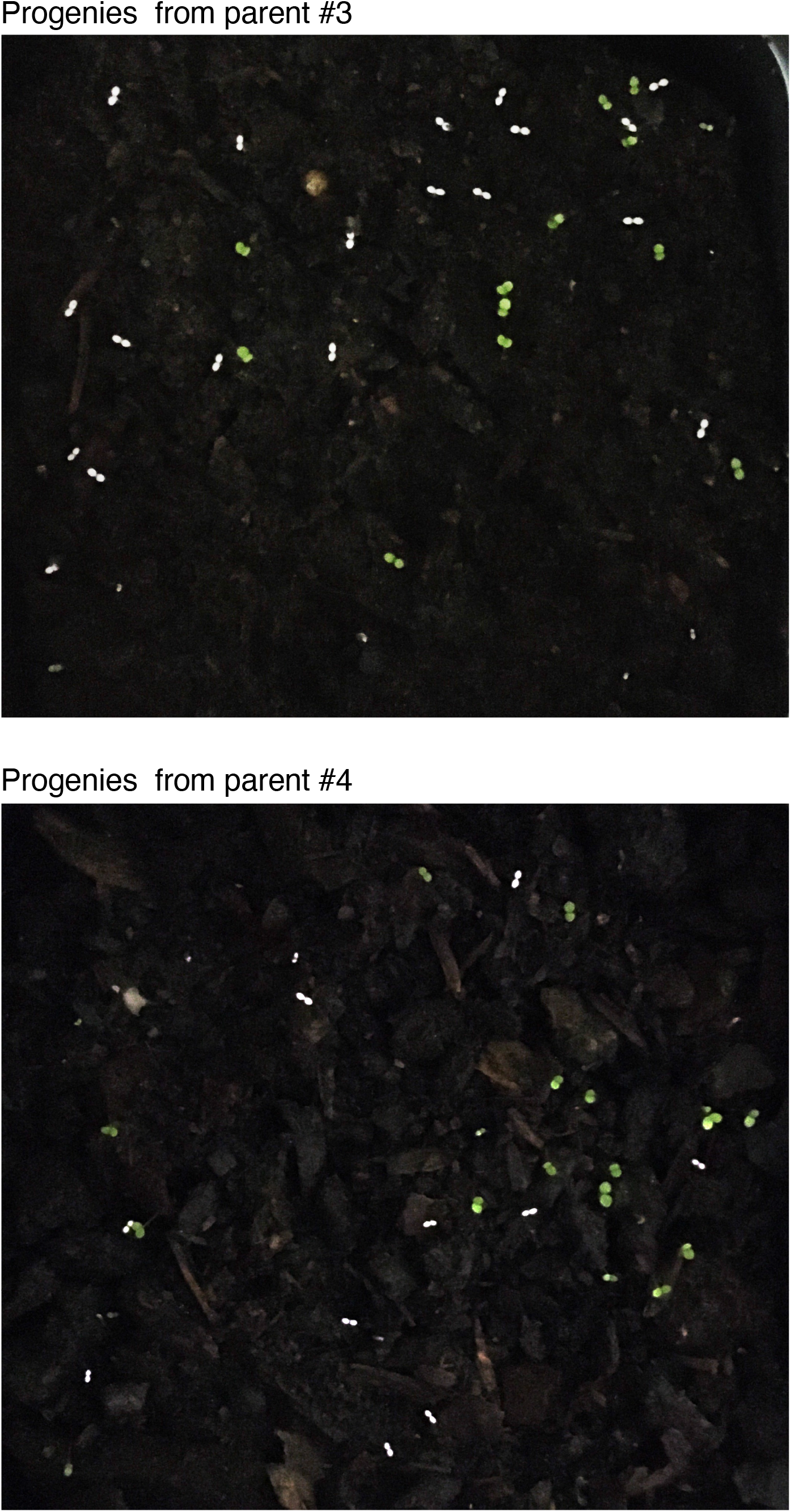
Heritable editing phenotype of *AtPDS3* in M1 progenies. M1 progenies from two parental plants infected with TRV1+TRV2::sgRNA^AtPDS3^-tRNA^Ileu^ showing a photobleaching phenotype.

**Supplementary Fig. 6.**
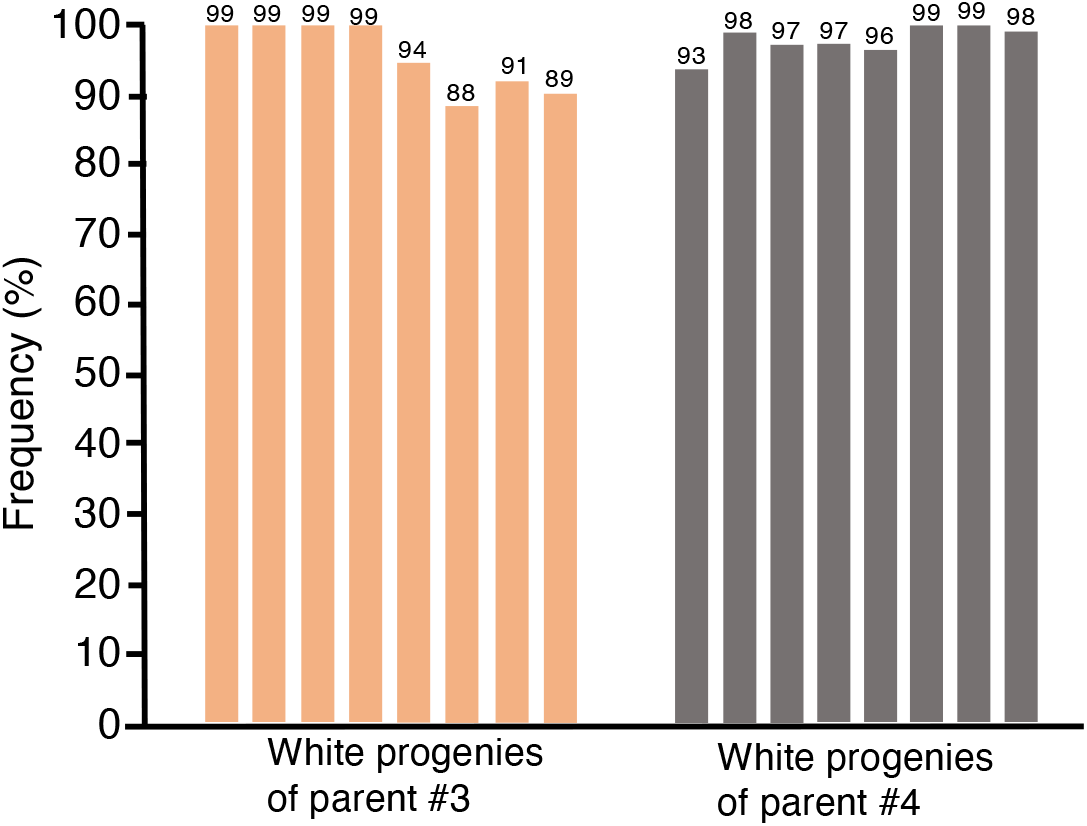
Indel mutation frequencies in *AtPDS3* in M1 progenies. Indel mutation frequencies assessed by Sanger sequencing of amplicons of *AtPDS3* target site followed by Synthego ICE analyses of M1 progenies of parental plants 3 and 4 infected with TRV1+TRV2::sgRNA^AtPDS3^-tRNA^Ileu^. Numbers shown on top of each bar indicate the percentage of indels.

**Supplementary Fig. 7.**
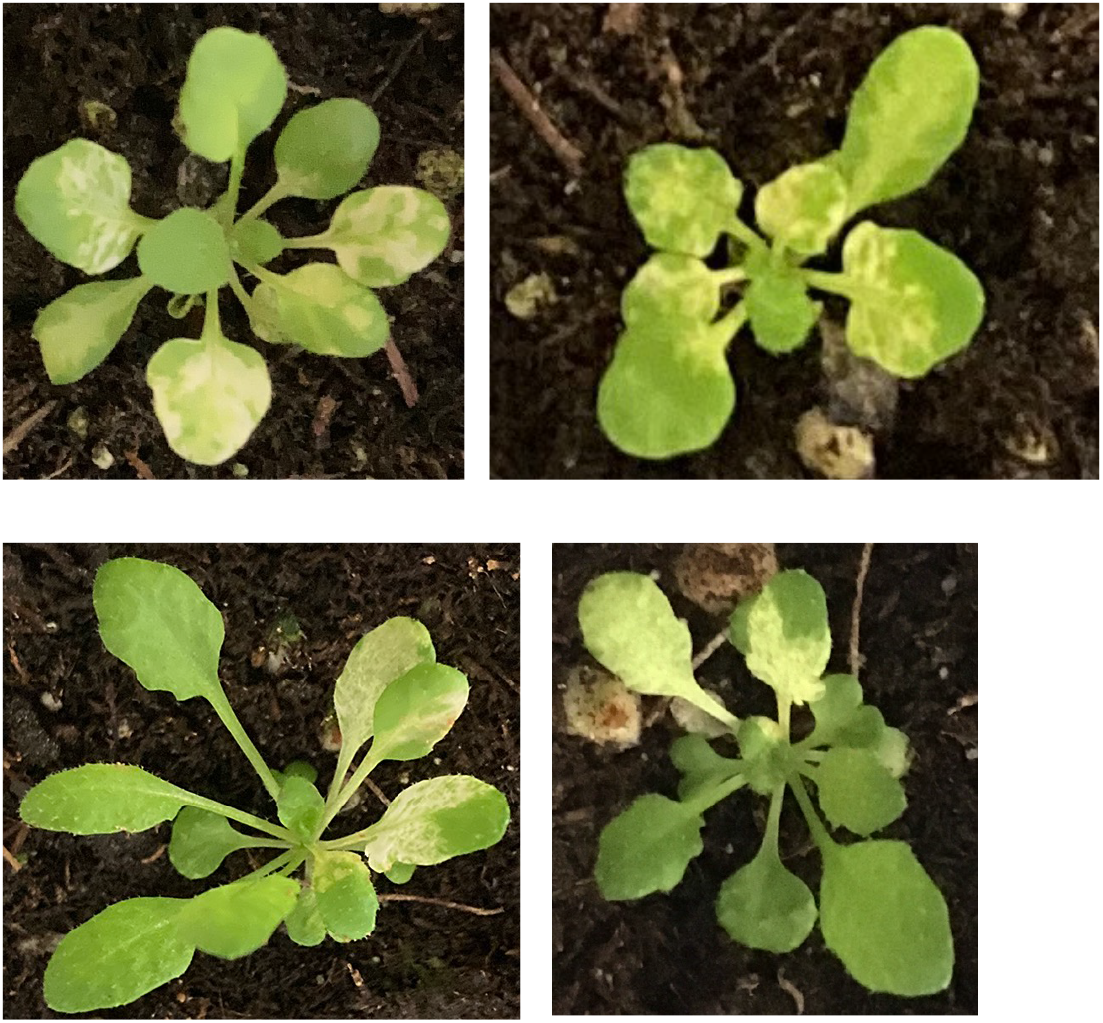
Plants showing *AtCHLI1* and *AtCHLI2* somatic editing phenotype.

**Supplementary Fig. 8.**
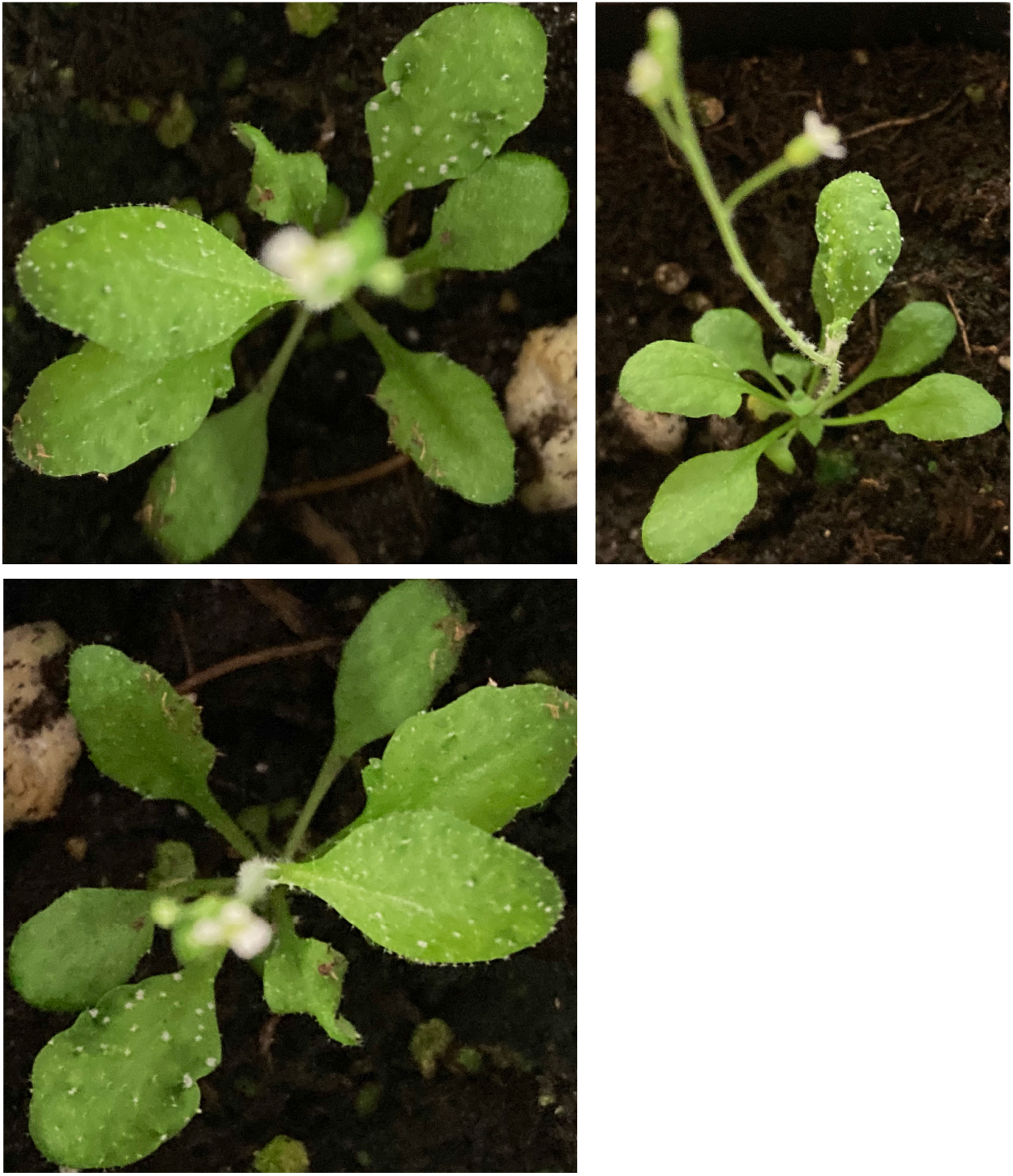
Plants showing *AtTRY* and *AtCPC* somatic editing phenotype.

**Supplementary Fig. 9.**
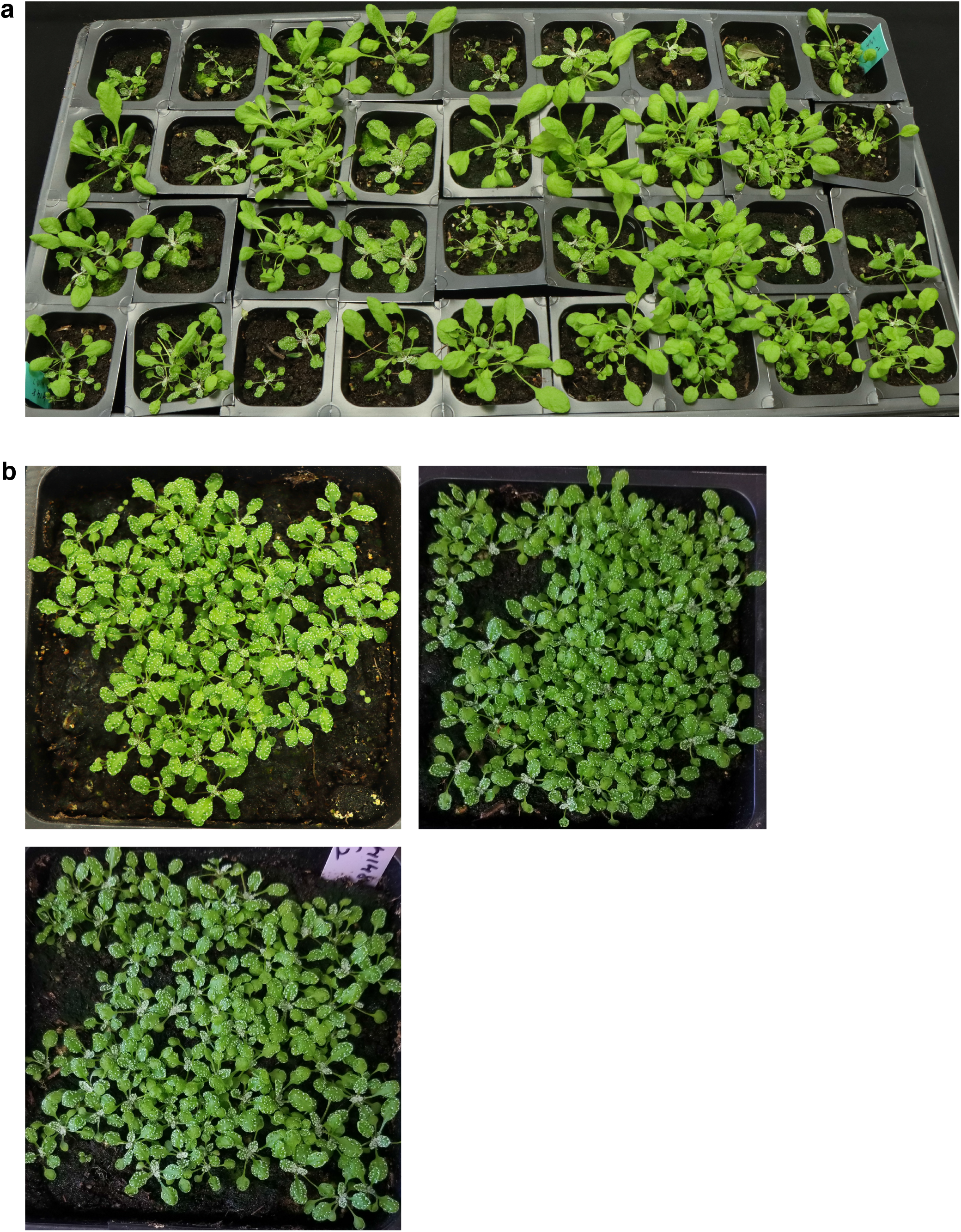
M1 and M2 progenies with leaf trichome phenotype due to editing in *AtTRY* and *AtCPC*. **a** M1 progenies from seed collected from a parental plant infected with TRV1+TRV2::sgRNA^AtTRY/AtCPC^-tRNA^Ileu^. **b** M2 progenies from seed collected from three independent M1 progenies.

## Notes

### Competing Interest Statement

The authors have declared no competing interest.

